# EFP Analyzer: A fast, accurate, and easy-to-teach program for analyzing Extracellular Field Potentials from iPSC-derived cardiomyocytes

**DOI:** 10.1101/2024.10.21.619481

**Authors:** Nidhi Patel, Alex Shen, Yuko Wada, Marcia Blair, Devyn Mitchell, Loren Vanags, Suah Woo, Matthew Ku, Kundivy Dauda, William Morris, Minjoo Yang, Björn C. Knollmann, Dan M. Roden, Joe-Elie Salem, Andrew M. Glazer, Brett M. Kroncke

## Abstract

**Rationale:** Induced pluripotent stem cell-derived cardiomyocytes (iPSC-CMs) are an emerging model for determining drug effects and modeling disease. Specialized devices can generate Extracellular Field Potential (EFP) measurements from these cells, analogous to the ventricular complex of the electrocardiogram.

**Objective:** The objective of this study was to develop an easy-to-use, easy-to-teach, reproducible software tool to measure EFPs.

**Methods-Results:** We present the EFP-Analyzer (EFPA), a semi-automized analyzer for EFP traces, which identifies and averages beats, identifies landmarks, and calculates intervals. We evaluated the tool in an analysis of 358 EFP traces from 22 patient-derived lines. We analyzed spontaneously beating iPSC-CMs, as well as optically paced iPSC-CMs through channelrhodopsin. We developed stringent quality criteria and measured EFP intervals, including Field Potential Duration (FPD). FPD from optically paced iPSC-CMs were shorter than those of spontaneously beating iPSC-CMs (283.7.0±54.2 vs. 293.0±47.5, p: 0.32, respectively). We further analyzed the usability and data replicability of EFPA through an inter-intra observer analysis. Correlation coefficient for inter-reader tangent and threshold measurements for these FPD ranged between r: 0.93-1.00. Bland-Altman plots comparing inter observer results for spontaneously beating and paced iPSC-CMs showed 95% limits of agreement (−13.6 to 19.4ms and −13.2 to 15.3ms, respectively). The EFP-analyzer could accurately detect FPD prolongation due to drug (moxifloxacin) or pathogenic loss of function mutations (*CACNA1C* N639T). This program is available for download at https://github.com/kroncke-lab/EFPA. The instructions will be available at the same listed website under the README section of the Github main page.

**Conclusions:** The EFP-Analyzer tool is a useful tool that enables the efficient use of iPSC-CMs as a model to study drug effects and disease.

## INTRODUCTION

Since their discovery in 2006,^1^ induced pluripotent stem cells (iPSCs) have become a transformative tool for modeling cell behavior, drug response, and disease.^2^ A number of groups have developed methods for differentiating iPSCs into cardiomyocytes (CMs).^3^ These induced pluripotent stem cell derived cardiomyocytes (iPSC-CMs) are an emerging model for the study of drug safety screening, in particular for detecting drugs that prolong the cardiac action potential duration.^4,5^ iPSC-CMs have also been used to study genetic background variability in electrocardiographic properties and drug response.^6^ In addition, gene-editing methods such as clustered regularly interspaced short palindromic repeats (CRISPR) allow further studies of mutation effects on iPSC-CMs.^7^ Advancements in optogenetic methods have allowed for the expression of channelrhodopsin, a light activated ion channel, in iPSC-CMs to enable optical pacing.^8^ More accurate investigations of cardiac function and pharmacological effects make it possible to precisely and non-invasively control cardiomyocyte activity. Although iPSC-CMs are thought to represent a relatively immature state of cardiac differentiation, recent methods have resulted in improved ventricular cardiomyocyte differentiation, successfully profiling all major cardiac currents in iPSC-CMs.^9^ The Extracellular Field Potential (EFP) measures the ion movement across cell membrane during contraction via adjacent electrodes.^10^ EFP signatures directly represent a cardiomyocyte’s beating pattern and could be considered analogous to the electrocardiogram (ECG). Previous literature defined the analogous time-based parameters as depolarization time (DT), repolarization time (RT), and maximum field potential duration (FPDmax).^11^

ECG analysis software automation is widespread with extensive validation regarding consistency of these measurements.^12^ The heart has large conduction distances and regions with distinct beating patterns, such as the atria, ventricles, and his-Purkinje complexes. In contrast, iPSC-CM monolayers are a relatively uniform substrate with small conduction distances. Although the EFP representation appears analogous to the ventricular complex (Depolarization and Repolarization) of the ECG (Figure 1), relatively little work has demonstrated the consistency or reliability of EFP measurements, including intra and inter observer reliability. In addition, no standardized analysis tools have been made to analyze EFP morphology and parameters. Previous studies have adapted ECG analyzers such as LabChart designed for *in vivo* animal analysis, used software built-in to the devices, or used custom analysis methods without assessing reliability of these strategies.^13–15^

**Figure 1.**
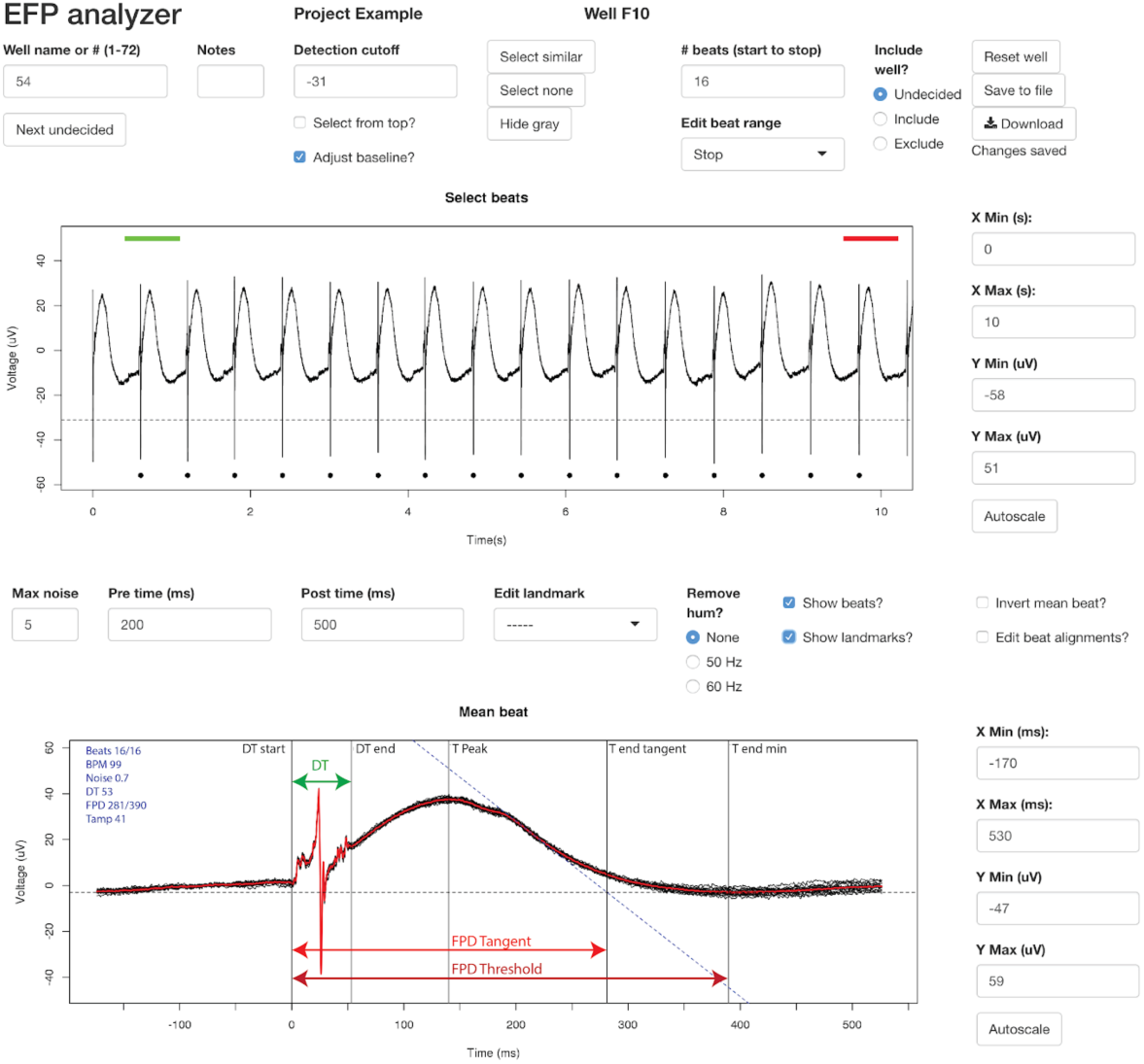
EFP Analyzer Software User Interfa

We present the EFP-Analyzer (EFPA), a semi-automated software tool for analysis of EFP traces. We use the EFP-Analyzer to analyze a large dataset of 358 control EFPs from 22 control patients and provide guidelines for stringent quality criteria. We show that this tool is fast to use, and generates consistent, reliable measurements. The EFP Analyzer can accurately detect changes in FPDmax from QT prolonging drug (moxifloxacin) or mutation (CACNA1C N639T). This tool will help facilitate the use of EFPs in iPSC-CMs for drug screening and disease modeling.

## Materials and Methods

### Generation of iPSC lines

Eight female and seven male patients with no known congenital arrhythmia syndromes were chosen (Figure S1). All patient-derived lines were generated with patient consent under Vanderbilt IRB #040551. Peripheral blood mononuclear cells were isolated from blood and reprogrammed into iPSCs using electroporation delivery (Neon system, ThermoFisher) of non-integrating episomal vectors (Epi5 Episomal iPSC Reprogramming Kit, ThermoFisher). iPSC-like colonies were manually moved to hES-grade matrigel (Corning) coated 24-well plates between day 14-30 post-electroporation. Individual colonies were expanded for validation and line maintenance. Selected iPSCs were validated by staining with OCT4 (Cell Signaling, #2750), SSEA4 (DSHB, MC-813-70), SSEA3 (Millipore, MAB4303), and Tra1-60 (Millipore, MAB4360). All iPSC lines used in this study were karyotype normal (Genetics Associates, Nashville, TN).

### Cardiomyocyte differentiation

IPSC’s were cultured in mTeSR media for 4-5 days to achieve ∼80% confluency. Media was changed to cardiac differentiation medium: RPMI 1640 (11875, Life Technologies), 2% B-27 minus insulin supplement (A1895601, Life Technologies), and 1% Penicillin-Streptomycin (Life Technologies). From days 0-2 the media was supplemented with 6 μM CHIR99021 (LC Laboratories), and from days 3-5 with 5 μM IWR-1 (Sigma). From days 13-19 metabolic selection medium was used: RPMI1640 without glucose (11879, Life Technologies), 2% B-27 minus insulin supplement (A1895601, Life Technologies), and 1% Penicillin-Streptomycin (Life Technologies). For days 19+ cardiomyocyte media was used. Media was exchanged every other day. Selected iPSC-CM were validated by staining with alpha-actinin (Sigma). Optically paced cells were treated from days 28-30 with 6500 genome copies/cell of AAV1.CAG.hChR2(H134R)-mCherry.WPRE.SV40 (Addgene20938M) purchased from the University of Pennsylvania Vector Core. At day 30 cells were replated into CardioExcyte96 sensor plates (Nanion) coated with 50 µl of 1:100 Matrigel, and cells were studied at day 38-42 post-differentiation.

### EFP measurement

EFP and impedance signals were measured on the CardioExcyte96 (Nanion) 8-12 days after replating (day 38-42 post-differentiation). Cells were kept at 37°C, 80% humidity, and 5% CO_2_. For optical pacing, a stimulating optical lid (Nanion) was used, which exposed all wells to 470nm wavelength light. Cells were exposed at 1 Hz to 5 msec-long pulses of light at 30 mA power. 20-30 second sweeps were recorded at an acquisition rate of 0.1 ms. Raw EFP and impedance measurements were converted into tab-delimited files and processed in the EFP-Analyzer tool.

### EFP-Analyzer analysis

A flowchart of the analysis logic is presented in Figure 1. The user is first presented with a data input screen (Figure 1) which allows the import of a tab delimited file containing time in seconds in column 1 and EFP measurements from multiple samples (wells) in columns 2+. Alternatively, the user can also load a previously analyzed project. Upon loading the input files and selecting initial settings, screen 2 is presented, which contains the main analysis. The top plot displays the entire sweep, from which beats can be selected automatically or manually for analysis. Beats are averaged and displayed in plot 2. Beats deviating from the mean beat by more than the noise cutoff are eliminated. Landmarks for the DT complex, the T peak, the *T_end_* tangent point, and the *T_end_* threshold point are selected automatically or manually (Figure 1). The user can navigate between samples and choose which samples to include or exclude. Measurements and plots can be downloaded to the user’s computer as a series of .csv and .pdf files. Additional EFP Analyzer features are described in the supplemental methods.

### Analysis of control EFPs

22 control lines were selected across 15 AAV-treated and 10 non-AAV treated lines (Figure 2). All EFPs were classified by morphology into one of 3 categories: standard, low amplitude, and biphasic (Figure 3). Low amplitude traces were defined as traces with a repolarization wave maximal amplitude (T peak) less than 15 uV. Biphasic traces were defined as those having two distinct repolarization waves in opposite directions, each later than 50 ms after the beginning of depolarization. Recordings were conducted in 20-30 second intervals at an acquisition rate of 0.1 ms. The mean beat was defined by the average of the beginning of the depolarization time to the end of the T wave from selected beats. Irregular traces were excluded, as standard, repeatable beats were preferred to avoid the complexity of median representative beat analysis. Optically paced traces not at 1 beat per second were excluded. Each EFP was then measured with the EFP Analyzer tool as described above.

**Figure 2.**
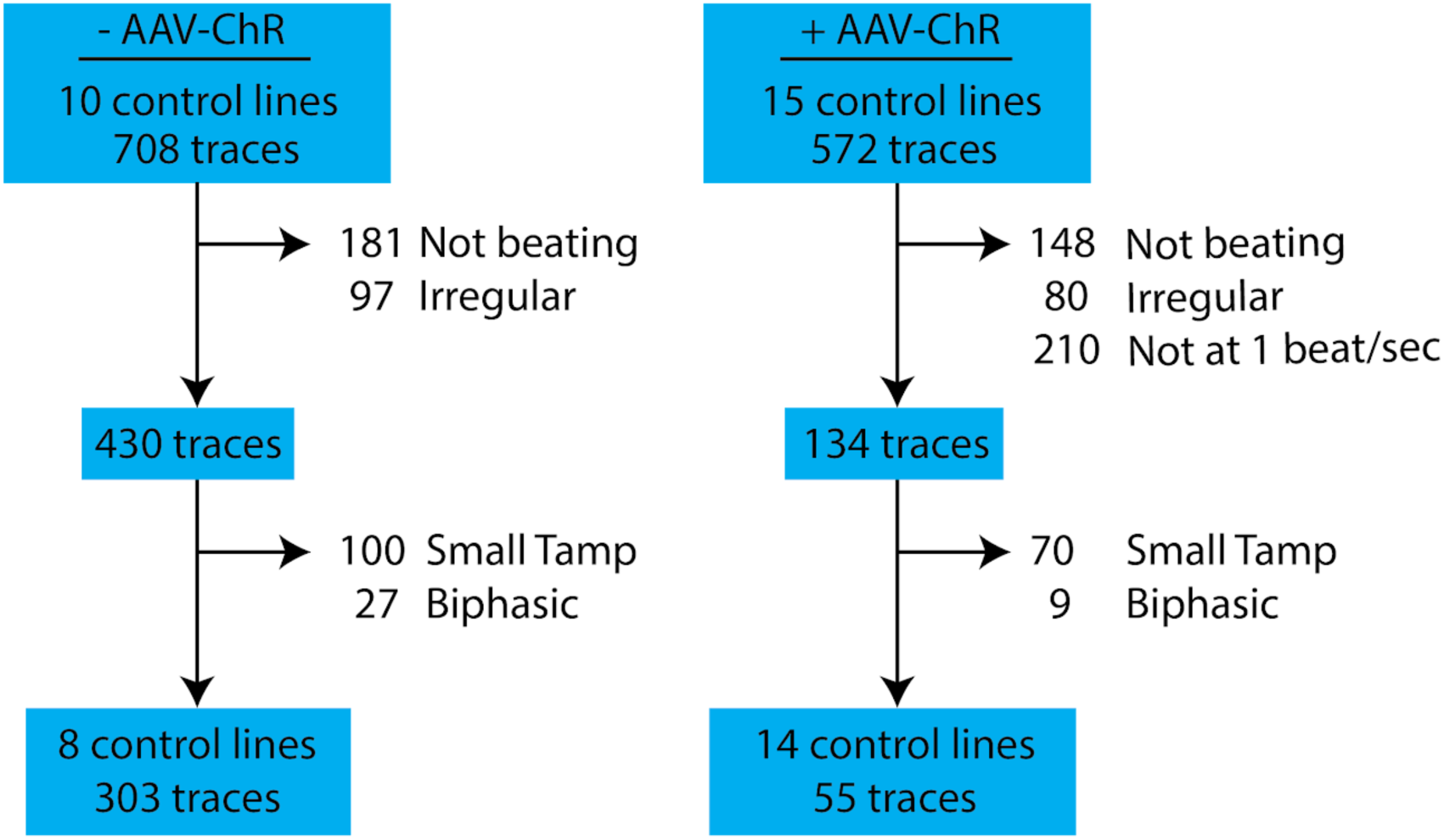
Flow Analysis Chart of Morphology between AAV-Chr & no AAV-Chr

**Figure 3.**
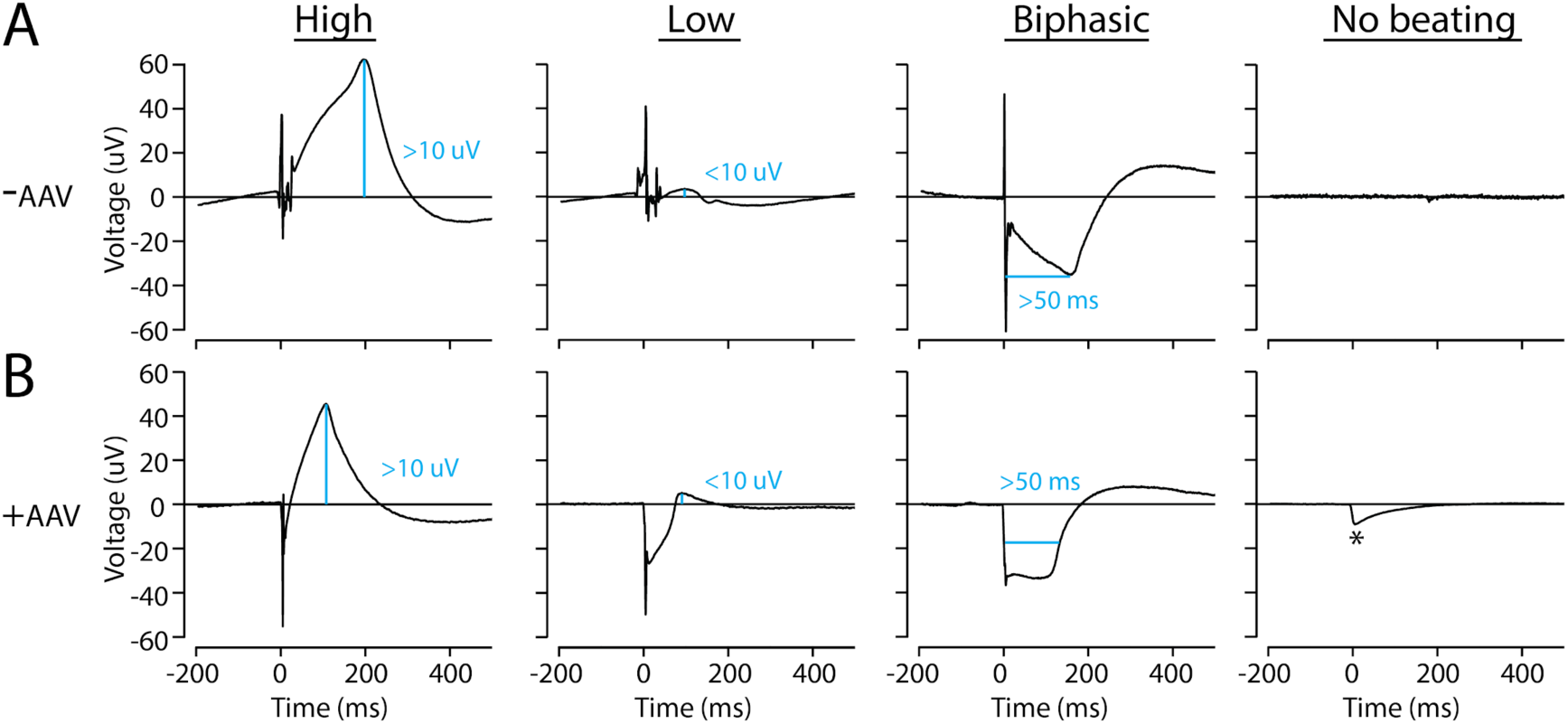
EFP Morphologies for Spontanously Beating and Paced Cells A) Example EFP morphologies for spontaneously beating cells (-AAV) B) Example EFP morphologies for paced cells (+AAV). * Indicates a light pacing artifact.

### Analysis of EFPs after exposure to an IKr-Blocker or to pathogenic genetic variant in CACNA1C

Moxifloxacin, a hERG channel blocker,^16^ was applied to patient generated iPSC-CMs without known congenital arrhythmogenic genetic background in increasing doses (10 uM, 30 uM, 100 uM, 300 uM) at 1-hour intervals. The iPSC lines heterozygous for CACNA1C N639T were generated as previously described.^11,17^ Statistical analyses were performed in R (version 4.1.2).

## RESULTS

### Description of EFP-Analyzer software tool

We developed the EFP-Analyzer software tool, a Shiny-based interactive tool, available for download at https://github.com/kroncke-lab/EFPA. The tool comprises two separate pages: a data load screen and a data analysis screen (Figure 1). In the data load screen, users upload a dataset of EFP signal versus time, formatted in a comma separated values (.csv) file, and select analysis parameters. In the data analysis screen, beats are selected and averaged, landmarks are placed and intervals are calculated. The philosophy of the program is that it is “semi-automated” and landmarks resemble to the one used to evaluate ventricular depolarization (QRS) and repolarization processes (QT) in human ECG. Therefore, these parameters are named DT and FPD (Figure 1). The program performs these tasks automatically once a new well is loaded. Users can modify the beat location and landmark placement.

### Analysis of 22 iPSC-CM lines

We applied the EFP-Analyzer to EFPs generated from the CardioExcyte96, a 96-well plate format EFP generator from Nanion. We evaluated iPSC-CM lines derived from individuals without apparent genetic arrhythmia disorders. From the recorded 25 patient lines, we screened a total of 1280 traces (30 seconds per trace; 8-150 traces per patient line). Some traces were analyzed with spontaneous beating, and some were treated with an AAV-ChR and optically paced. From an initial dataset of 708 spontaneous and 572 optically paced traces, wells that were not beating or irregularly beating were excluded leaving 303 spontaneous and 55 AAV traces from 22 patient lines (Figure 2). We determined the consistency of the EFP-Analyzer to measure FPDmax on EFP traces as a function of EFP morphology classified into 3 categories (standard, low amplitude, biphasic) (Figure 3). A random selection of 260 paced and 430 unpaced traces (stratified by EFP repolarization morphology) were repeated in measurement by an expert (AG), and by a novice (YW) who had a training session of 10 minutes in how to analyze the data and use the tool (∼10 EFP test measurements) (Figure 4). The consistency and bias, the mean difference between repeat measurements as a function of morphology traces are displayed in Table 2 and Table 3. The EFP with a standard morphology had the highest level of measurements reproducibility while EFP with low amplitude or biphasic morphology were poorly reproducible and highly scattered (Table 1).

**Figure 4.**
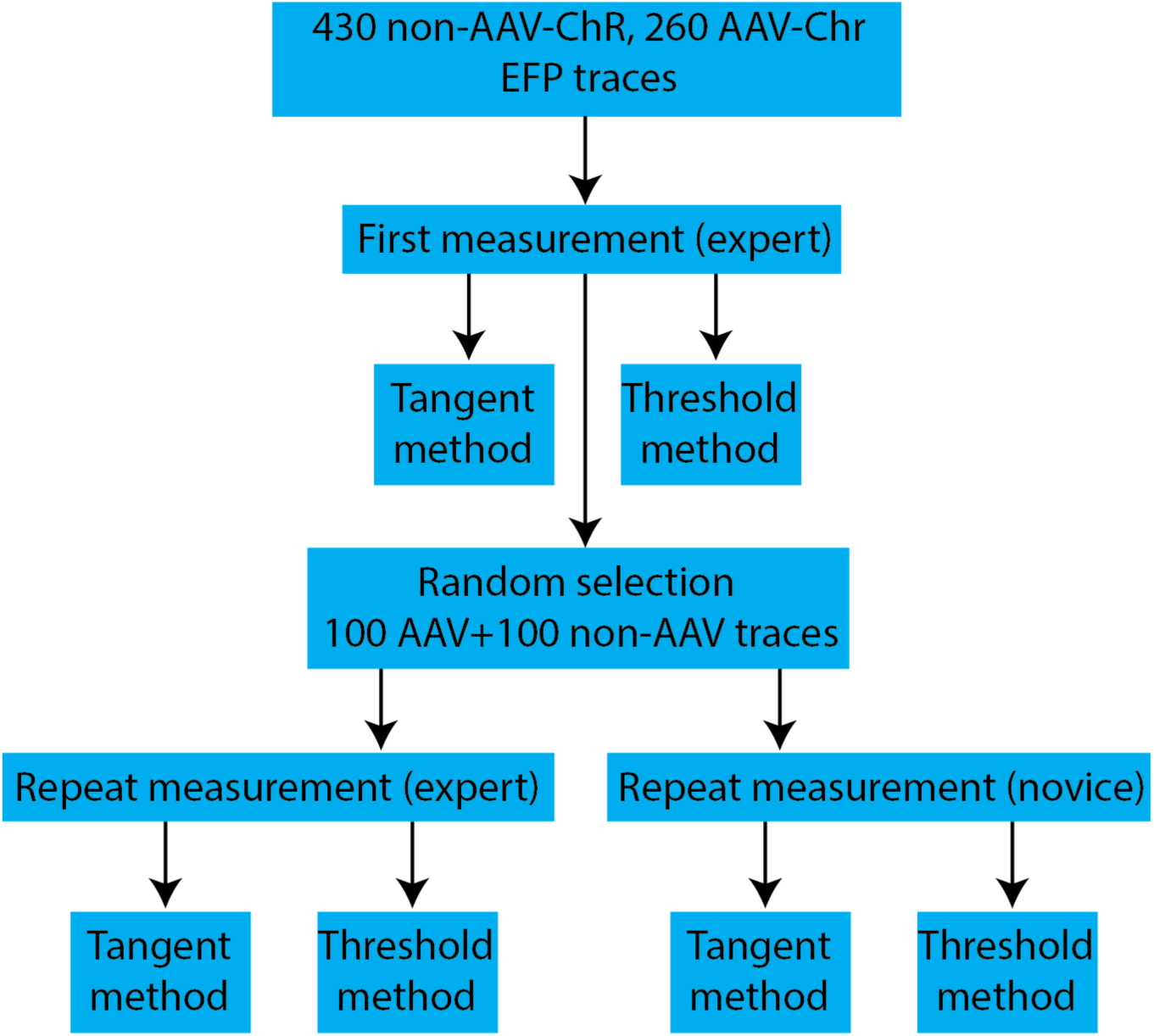
Flow Analysis Chart of Inter and Intra-Reader Variability Assessment

**Table 1.** Comparison of EFP parameters based on morphology & measurement method.

| AAV-ChR | Morphology | Number of wells | RR (ms) | EFP-DT | FPD tangent | FPD threshold | FPD tangent vs. threshold p-values |
| --- | --- | --- | --- | --- | --- | --- | --- |
| No | Standard | 303 | 798.6(314.3) | 58.9 (21.6) | 293.0 (47.5) | 461.0 (82.1) | <0.001 |
|  | Small Tamp | 100 | 778.0(202.2) | 59.8 (17.9) | 466.3 (170.6) | 603.5 (174.2) | 0.06 |
|  | Biphasic | 27 | 839.1(512.4) | 108.5 (123.5) | 602.9 (280.4) | 736.1(349.3) | 0.96 |
| Yes | Standard | 55 | 995.4(4.99) | 26.4 (14.7) | 283.7 (54.2) | 442.9(109.2) | <0.001 |
|  | Small Tamp | 21 | 993.7(4.93) | 29.8 (28.1) | 542.8(182.2) | 690.2 (185.9) | 0.01 |
|  | Biphasic | 2 | 994.7(6.65) | 45.2 (26.7) | 369.9 (120.3) | 690.3 (199.1) | 0.22 |
| AAV-ChR (Yes vs. No) p-values | Standard | NA | <0.001 | <0.001 | 0.32 | 0.31 | NA |
|  | Small Tamp |  | <0.001 | <0.001 | 0.16 | 0.02 |  |
|  | Biphasic |  | 0.42 | 0.91 | 0.14 | 0.69 |  |

The numbers represent mean (standard deviation). *Abbreviations*: AAV: adeno-associated virus; EFP: extracellular field potential,
iPSC-CM: induced pluripotent stem-cell derived cardiomyocyte, CRISPR: clustered regularly interspaced short palindromic repeats;
FPD: Field Potential Duration; RR: time between two successive waves from the start of DT.

**Table 2.** Comparison of EFP consistency based on inter and intra reader and measurement method.

| AAV-ChR | Morphology | Number of wells analyzed | FPD tangent consistency intra-reader (ms) | FPD tangent consistency inter-reader (ms) | FPD threshold consistency intra-reader (ms) | FPD threshold consistency inter-reader (ms) |
| --- | --- | --- | --- | --- | --- | --- |
| No | Standard | 68 | -0.6 (11.2) | 2.3(13.1) | -3.6(21.3) | 3.9(23.4) |
|  | Small Tamp | 24 | -2.0 (15.9) | -2.8 (47.1) | -9.1(25.5) | 19.5(28.8) |
|  | Biphasic | 8 | -18.7 (64.9) | -39.2 (104.5) | -32.9(39.6) | 30.6(126.1) |
| Yes | Standard | 73 | -0.7 (8.6) | 1.0 (7.3) | -0.1(5.6) | -0.2(10.8) |
|  | Small Tamp | 25 | -5.2 (36.0) | -6.5 (43.3) | 0.7(5.5) | -7.3(34.3) |
|  | Biphasic | 2 | -2.4 (27.4) | 8.5 (12.0) | -31.2(44.1) | -31.2(44.1) |

Abbreviations: ChR: channelrhodopsin

**Table 3.**
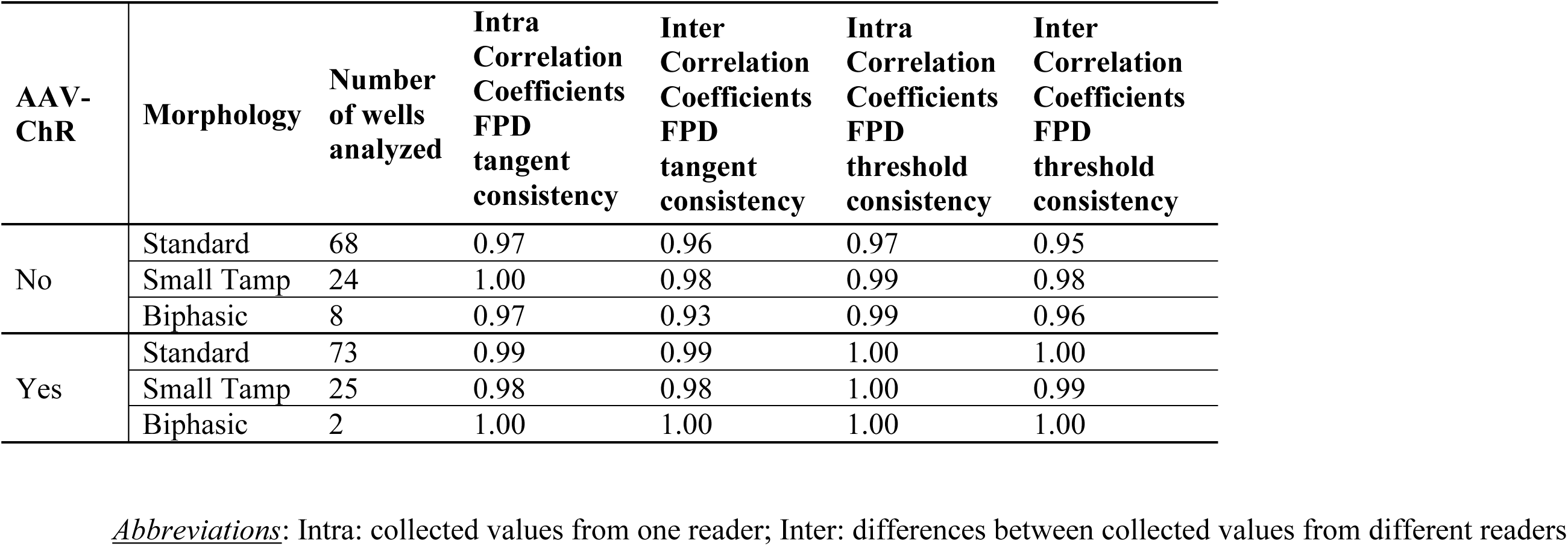
Comparison of FPD consistency based on inter and intra reader and measurement method

Therefore, we focused for the subsequent analyses only on the EFPs with standard trace morphology, 303 (303/708, 43%) spontaneous and 55 (55/572, 9.6%) high-quality AAV-ChR traces which compose the main dataset. All traces were measured by an expert analyzer (AG). Two methods were used, a simple threshold method and a tangent method, based on the different DT, reflective of measurement techniques used in human ECGs (Table 1). FPD assessment by tangent method was shorter and carried a lower dispersion than the FPD assessed by threshold method (283.7±54.2 vs. 442.9±109.2, p:<0.001, respectively, Table 1). Among the FPD (tangent method) measured by two readers, Bland-Altman plots results showed 95% limits of agreement ranging from –13.6 to 19.4 ms and from -13.2 to 15.3 ms for spontaneously beating and paced iPSC-CMs, respectively (Figure 5 and Table 2). Among the FPD (tangent method) measured by the intra reader, Bland-Altman plots results showed 95% limits of agreement ranging from -22.6 to 21.5 ms and from -17.4 to 17.0 ms for spontaneously beating and paced iPSC-CMs, respectively (Figure 5 and Table 2). Correlation coefficient for inter reader measurements for these FPD (tangent method) ranged between r:0.93-1.00 (Table 3). EFP-FDP (tangent method) were shorter in the group of paced EFP vs. spontaneous EFP (283.7±54.2 vs. 293.0±47.5, respectively) partly due to shorter EFP-DT in the group of paced EFP vs. spontaneous EFP (26.4±14.7 vs. 58.9±21.6, respectively). Data concerning Bland-Altman plots and intraclass correlation coefficient for intra-and inter-reader for EFP-FPDmax measured by threshold method are provided in Table 2. Data concerning differences between spontaneous and paced EFP for FPD measured by threshold method shared the same trend that with FPD measured by tangent method and are displayed in Table 1. Amongst the spontaneously beating and paced cells, FPD was not significantly correlated with RR (spearman rho= 0.183, p = 0.135; and rho = -0.165, p = 0.147; respectively, Figure S2).

**Figure 5.**
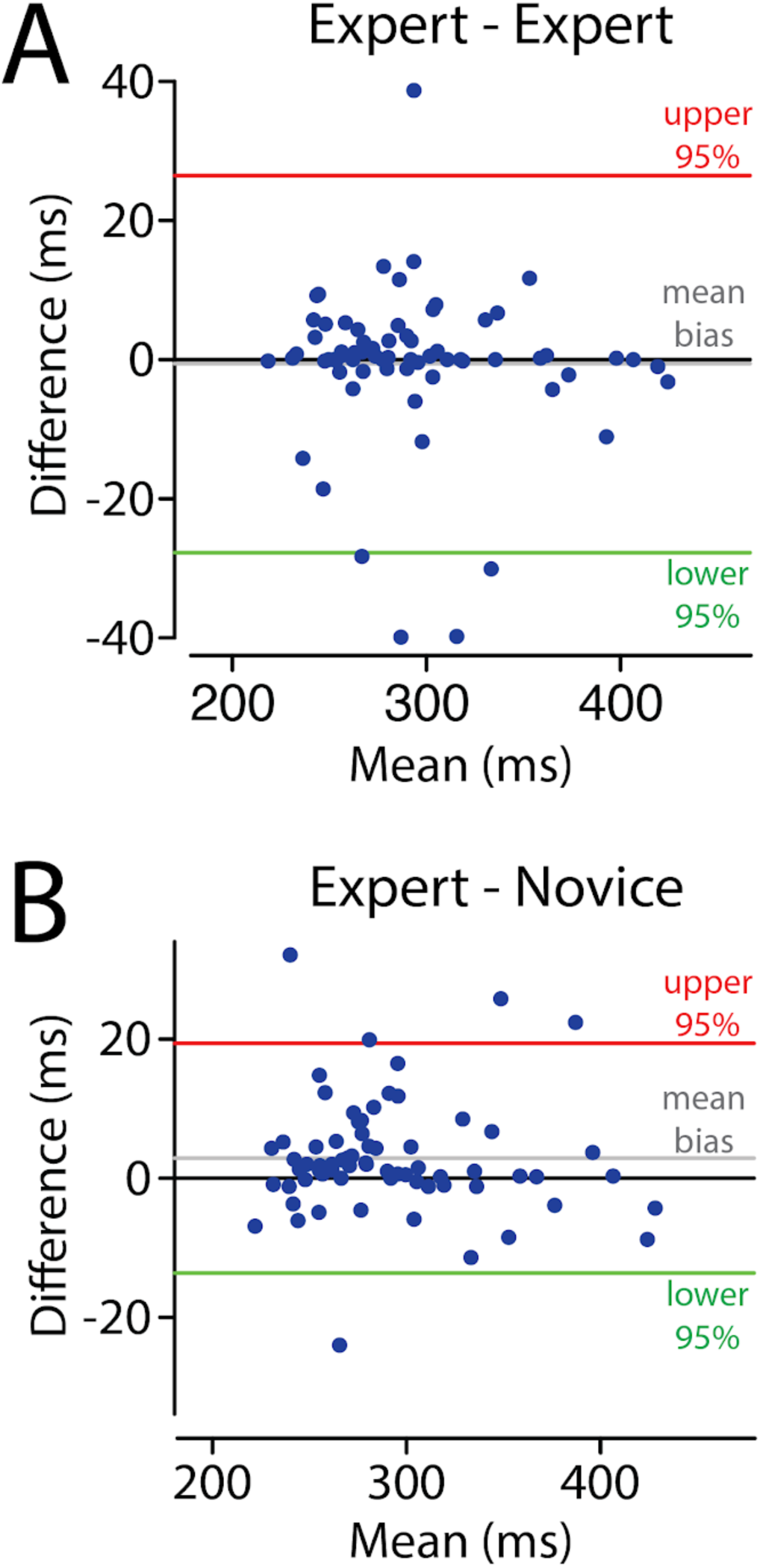
Comparison in FPD (ms) between Intra and Inter Reader A) Difference in FPD (ms) between intra-reader (expert-expert) B) Difference in FPD (ms) between inter-reader (expert-novice)

### Application to drug response and disease

We analyzed the EFP-Analyzer tool’s ability to identify changes to EFP pattern/landmarks in response to drugs and in the case of individuals with disease. We first analyzed the action potential-prolonging drug moxifloxacin. We successfully detected an increase in FPDmax in response to increasing doses of moxifloxacin, with 119 ± 9.9 ms prolongation at 100 uM (Figure 6). Next, we re-analyzed a previously published cell line carrying a pathogenic Long QT Type 8 *CACNA1C* variant N639T. Consistent with the previous study, relative to isogenic controls (non-CRIPSR edited) the mutation positive cell line had FPDmax prolongation in optically paced cells, (n = 15 cells/line, p < 0.002, two-tailed t test, Figure 6).

**Figure 6.**
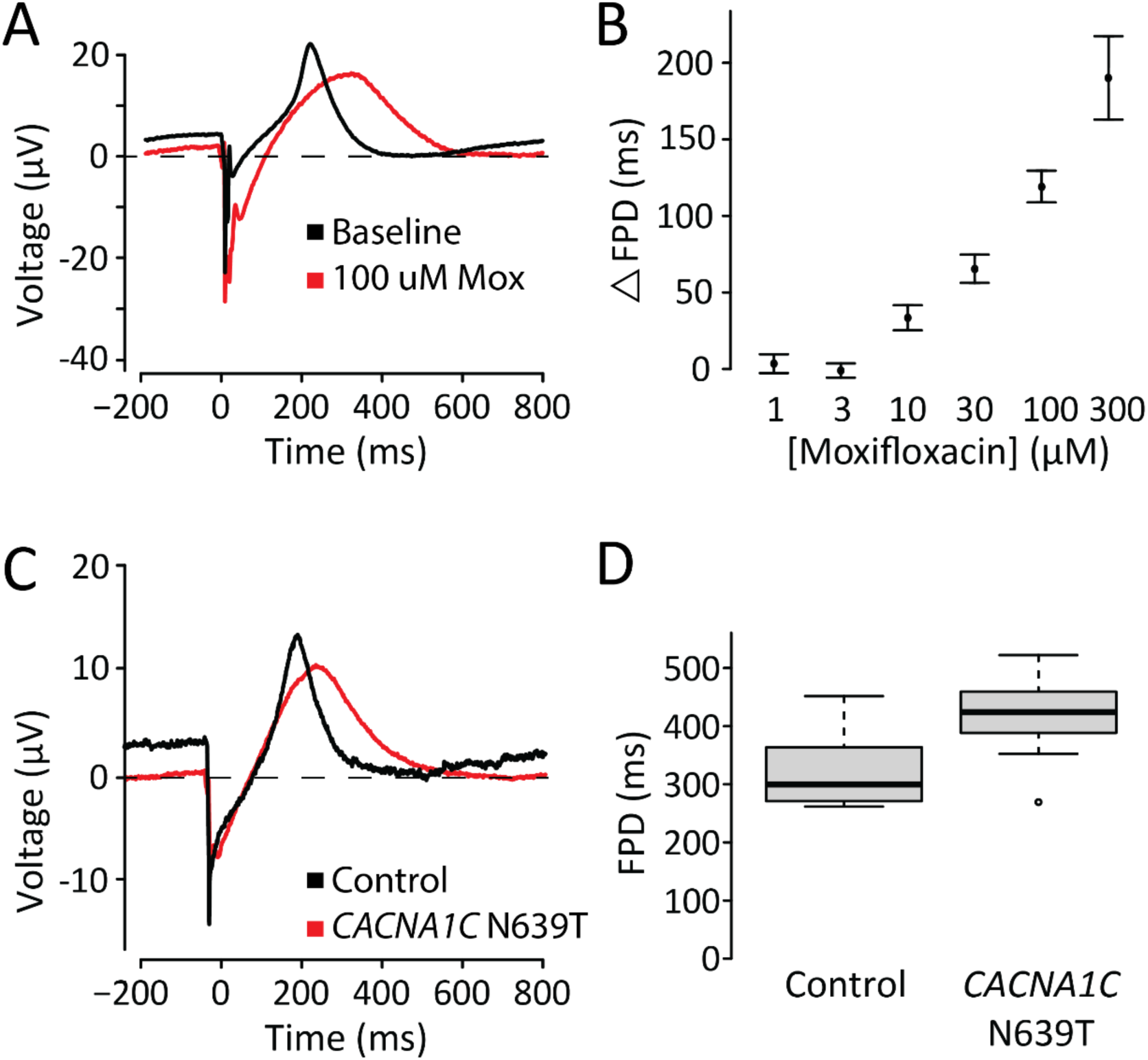
FPD response in an exposure to IKr Blocker and CACNA1C N639T A) Representative EFP trace from an example cell treated with 100uM Moxifloxacin. B) As the doses of Moxifloxacin increased, the change in QT was measured. Each measurement represents n= 11 cells. C) Representative EFP trace from an example N639T mutation positive cell. D) The mutation positive cell line had prolongation in optically paced cells, (n = 15) cells expressing CACNA1C N639T mutation exhibit a prolonged FPD compared to control cells (n = 19).

## DISCUSSION

### Summary

The EFP-Analyzer (EFPA) aims to streamline cardiac disease modeling. We showed consistency in measurements in interpretation of beats and landmarks across 22 patient lines with 358 traces. We were able to reliably detect pharmacological (moxifloxacin) and pathological alterations (CACNA1C N639T) from functional signatures. We further established strong inter-reader correlation coefficients (r:0.93-1.00) and good agreement in Bland-Altman. This technology not only allows for high-quality data capture, but it also ensures replicability across multiple users and studies^10^, representing a significant advancement in integrating iPSC-CMs in drug screening paradigm. EFPA’s versatility and convenience of use are critical for promoting wider adoption in academic and pharmaceutical research, notably in improving the efficiency, accuracy, and accessibility of EFP measurements.

### The EFP Analyzer is a fast, consistent tool for data analysis

The data derived from EFPA is robust and consistent between multiple observers. Analysis of 9 replicate sweeps in a 96 well plate was 12 ± 3 minutes. The mean difference between observers was 20-25 ms for unpaced and 8-12 ms for paced cells. This approaches the robustness achievable in an ECG’s QTc, typically considered to be around 11.8 ms.^12^ We speculate that uniform synchronous depolarization of an entire well using optical pacing resulted in a more repeatable (and 25 ms shorter) FPDmax than spontaneous cell activation in a well, which requires the signal to spread throughout the well from a point of activation.

### Recommendations for improving EFP quality/robustness

We observed a range of EFP morphologies, which we termed standard, low amplitude, and biphasic. We found that according to a variety of metrics, low amplitude and biphasic EFPs generated non-robust data, leading to highly variable, inconsistently measured values. We therefore recommend that these classes be removed. The tangent approach also improved the robustness of the EFPs according to these metrics. We note that these parameters were optimized on EFPs generated from one device, the Nanion CardioExcyte 96. Alternative cardiac screening instruments may differ in system defaults such as sampling rates, leading to potential differences in results.

For ECG analysis, a return to baseline of the QT is commonly used with multiple input convergence. However, in some cases the T wave has an extended unclear return to baseline particularly the lower the T peak is in term of amplitude, and a tangent method (drawn at the steepest point) yields more robust FPD measurements.^18^ We observed that some EFPs had long extended repolarization waves, analogous to T waves in the ECG. Thus, we employ the tangent method and observed a decrease in the standard deviation of the tangent and threshold method across many wells, which suggests both approaches are accurate/repeatable; we similarly observed more robust measurements when comparing across multiple observers.

### Comparison to other methods/tools

The CardioExcyte 96 has well been defined by several groups in high throughput genomic and pharmacological investigations and has since proven to be invaluable across disciplines^3,17,19–21^. It has built-in software to calculate intervals, however we found that the EFP Analyzer was a better fit for analyzing the data flexibly and storing and managing the data. The EFP Analyzer has increased capabilities, such as the tangent method, an option to manually adjust the beat locations, and persistent data storage. Additionally, the EFP analyzer standardizes the baseline start for each complex and allows for additional analyses such as calculating T-Amp, therefore increasing the accuracy of evaluations. Another advantage of the EFP Analyzer is that it is semi-automated; for many traces it correctly identifies all the landmarks without any manual changes. Other EFPs with complicated/unclear patterns or with high background/artifact need manual intervention to optimize. Another advantage is that it also functions as a data organizer that saves data which simplifies analysis, reviewing, and editing steps for collaborating researchers.

### Limitations

The ability of this software to analyze EFP repolarization wave characteristics was demonstrated in our investigation. Even though T peak amplitude and T peak T_$%&_ are important clinical metrics, we did not test the capability to analyze these characteristics.^12^ Furthermore, we did not study HR dependence of field potential duration. We observed a RT-peak amplitude decrease with moxifloxacin incubation, which calls for further investigation. Additionally, this method has also been compatible with *CACNA1C* N639T, a cell line with a congenital Long QT syndrome type 8. This program was customized for Nanion CardioExcyte 96 device and datasets with fundamentally similar EFP shapes. Data processing devices with differing EFP characteristic output, such as MEAs, may pose as a limitation. Future research is needed to test this and discover the software’s maximum capabilities. We encourage software limit testing to discover maximum capabilities. All comments, suggestions, and feedback for EFP-Analyzer can be submitted on https://github.com/kroncke-lab/EFPA/issues.

## Funding

This research was funded by the National Institutes of Health: R01HL160863 (BMK), R00HG010904 (AMG), and R35GM150465 (AMG); and by the Leducq Transatlantic Network of Excellence Program 18CVD05 (BMK)

## Conflict of Interest

The authors declared no competing interests for this work

---

**Supplementary Figure 1.**
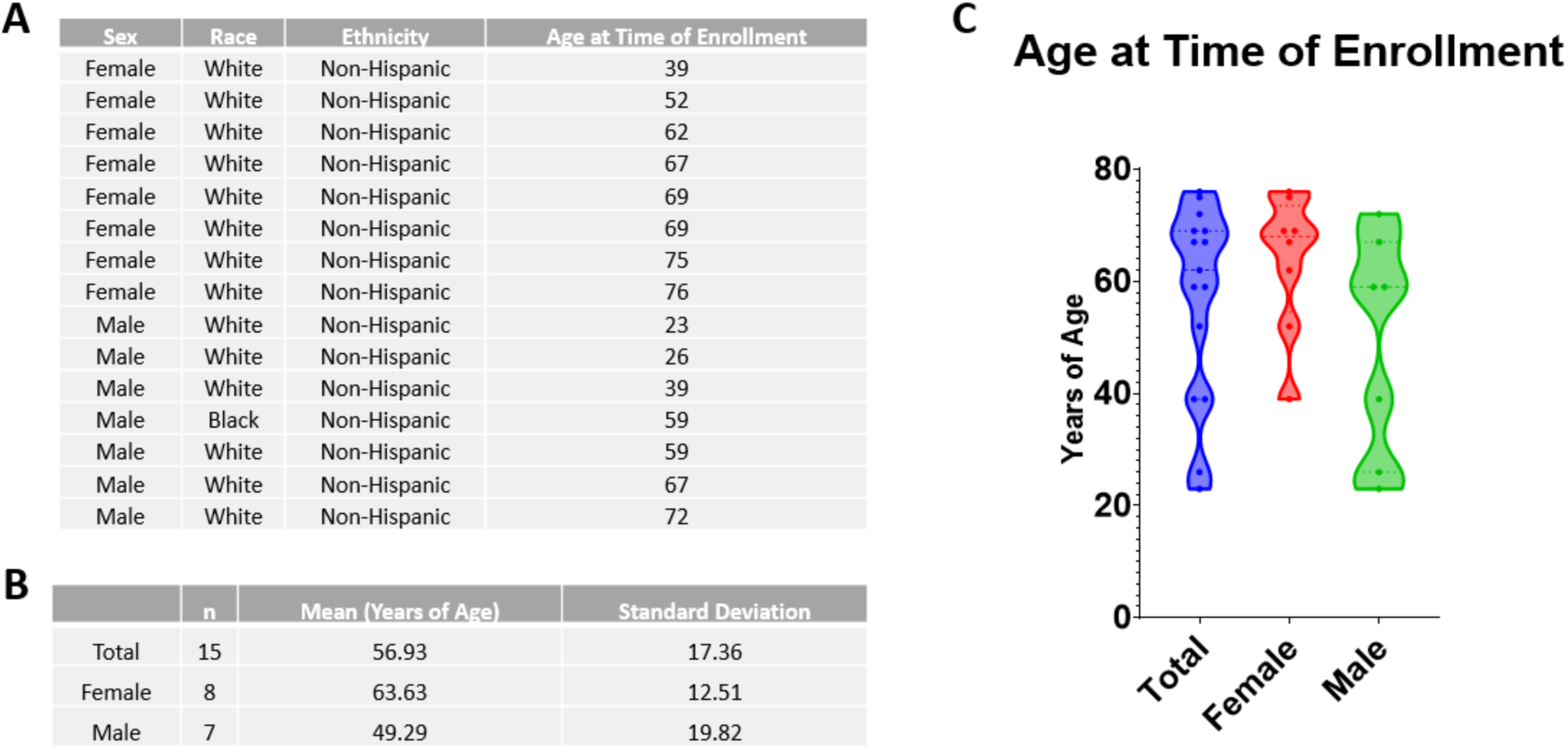
Patient Demographic Information A) Enrolled patient demographic: sex, race, ethnicity, and age of enrollment B) Mean patient age showing total n=15, female n=8, and male n=7 C) Volin plot depicting patient age at time of enrollment

**Supplementary Figure 2.**
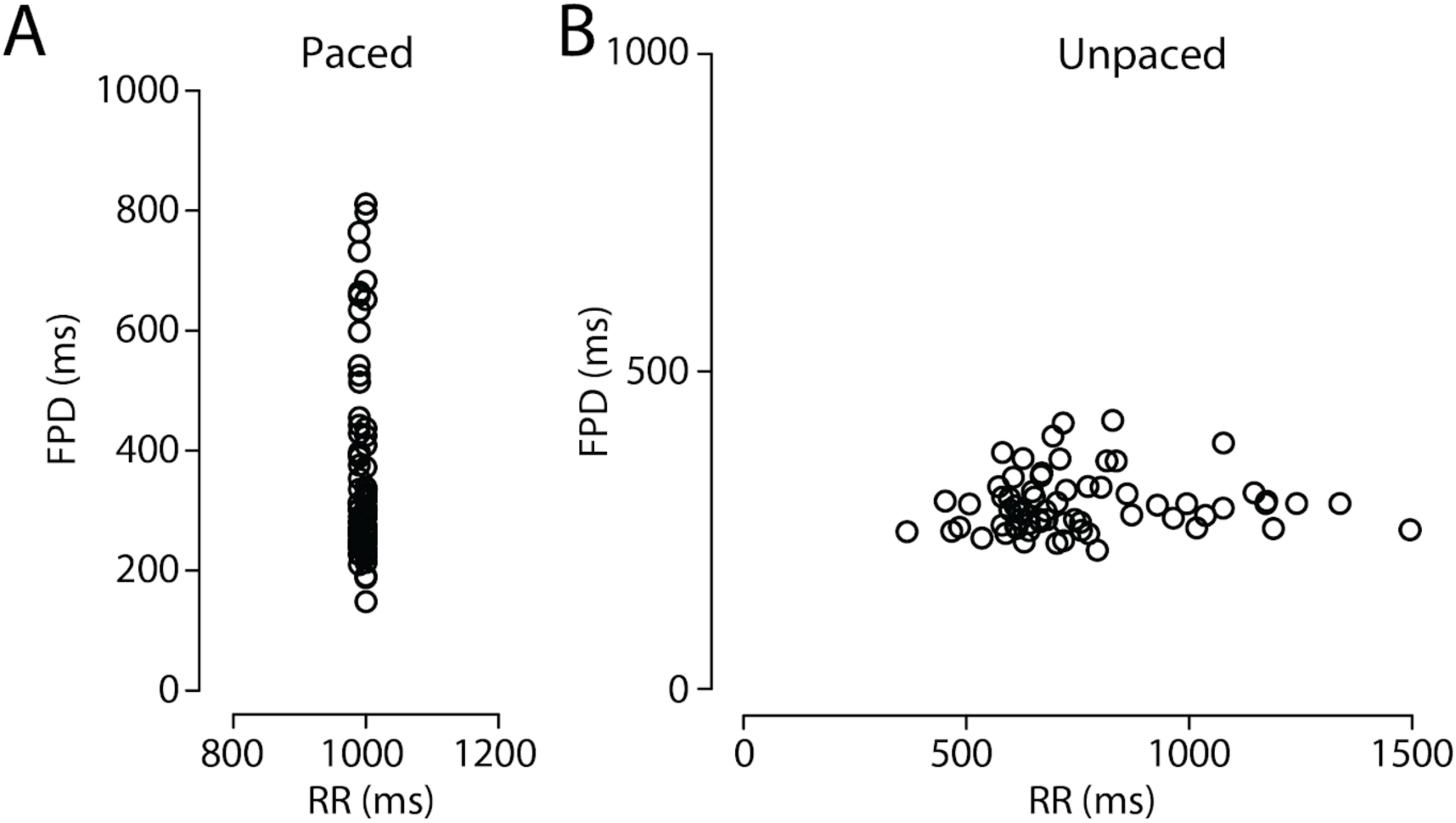
Comparison of RR and Field Potential Duration for Paced and Unpaced Cells A) Paced cells spearman’s rank correlation rho : S = 92171, p-value = 0.1474, rho = -0.165 B) Unpaced cells spearman’s rank correlation rho : S = 42805, p-value = 0.1352, rho= 0.183

## Notes

https://github.com/kroncke-lab/EFPA

### Competing Interest Statement

The authors have declared no competing interest.

### Summary of Updates

The paper format was edited and revised for clarity and readability.

